# Characterizing interactions between the microtubule-binding protein CLIP-170 and F-actin

**DOI:** 10.1101/2021.04.27.441644

**Authors:** Yueh-Fu O. Wu, Rachel A. Miller, Emily O. Alberico, Nora T. Nelson, Erin M. Jonasson, Holly V. Goodson

## Abstract

The cooperation between the actin and microtubule (MT) cytoskeletons is important for cellular processes such as cell migration and muscle cell development. Full understanding of how this cooperation occurs has yet to be sufficiently developed. The MT plus-end tracking protein (+TIP) CLIP-170 has been implicated in this actin-MT coordination by associating with the actin-binding signaling protein IQGAP1, and by promoting actin polymerization through binding with formins. Thus far, CLIP-170’s interactions with actin were assumed to be indirect. Here, we demonstrate that CLIP-170 can bind to filamentous actin (F-actin) directly. The affinity is relatively weak, but is strong enough to be significant in the actin-rich cortex, where actin concentrations can be extremely high. Using CLIP-170 fragments and mutants, we show that the direct CLIP-170:actin interaction is independent of the FEED domain, the region that mediates formin-dependent actin polymerization, and that the CLIP-170 F-actin-binding region overlaps with the MT-binding region. Consistent with these observations, *in vitro* competition assays indicate that CLIP-170:F-actin and CLIP-170:MT interactions are mutually exclusive. Taken together, these observations lead us to speculate that direct CLIP-170:F-actin interactions may function to reduce the stability of MTs in actin-rich regions of the cell, as previously proposed for EB1.

## Introduction

Components of the cytoskeleton are often described as having apparently independent localizations and activities. For example, actin accumulates at the cell cortex to maintain cell shape and promote whole-cell locomotion (1); MTs radiate from the center of typical animal cells to direct the position of cell organelles and promote intracellular transport (2). In addition to these seemingly individual activities, components of the cytoskeleton cooperate with each other to perform more complex cellular functions. The coordination and integration of the actin and MT cytoskeletons are known collectively as actin-MT crosstalk and are important for cellular processes such as cell division, establishment of cell polarity, neuronal regeneration, wound-healing, and muscle cell development (reviewed in (3–5)). However, while the significance of this crosstalk is clear, the mechanisms by which it occurs have yet to be fully elucidated.

Many mechanisms of actin-MT crosstalk have been studied, and they can be roughly categorized into three groups. First, shared signaling cascades can regulate the dynamics of both the actin and MT cytoskeletons (3–5). For example, Rho GTPases promote formin-dependent actin polymerization while also increasing stabilized microtubules near the leading edge of the cell (6). Second, crosslinking proteins can bridge the two cytoskeletons (3–5). This connection of the two filamentous systems can go through one or multiple proteins. For example, spectraplakins physically interact with both actin and MTs simultaneously (7), while the MT-binding protein EB3 connects the two cytoskeletal systems by binding actin through the actin-associated protein drebrin (8). Third, regulators of one filament type can bind or even be regulated by components of the other filament network (3)–5)). Examples include the observation that the actin nucleator formin binds and stabilizes MTs *in vivo* (9), and that MT plus-end tracking proteins (+TIPs) regulate actin polymerization as described below.

+TIPs are a subset of microtubule-associated proteins (MAPs) that track and regulate the dynamics of MT plus-ends. Accumulating evidence implicates +TIPs as mediators of actin-MT crosstalk (reviewed in (3-5)). For example, the key +TIPs EB1 and APC interact with formins, and this interaction is believed to regulate actin-MT coordination (9,10). In addition, APC directly binds to actin both *in vitro* and *in vivo* (11), and the C-terminal domain of APC promotes actin assembly (12,13); interestingly, binding of EB1 to APC downregulates APC-mediated actin assembly (14).

CLIP-170 was the first +TIP characterized (15,16); it regulates MT dynamics, promotes organelle-MT interactions (15,17), and binds to the core +TIP EB1 (18,19). Although the role of CLIP-170 in MT dynamics has been extensively-studied, the role(s) of CLIP-170 in regulating actin are still under investigation. Previous studies have shown that CLIP-170 regulates the actin cytoskeleton by two different mechanisms: (1) Rac1/Cdc42/IQGAP1 forms a complex with CLIP-170 to connect the MT and actin networks for mediating cell polarization and migration (20,21); (2) CLIP-170 promotes actin-polymerization through formins (22,23). These two mechanisms are relatively well-studied, but in both cases, the interactions between CLIP-170 and actin are indirect. We were interested in the possibility that CLIP-170 might bind to actin directly.

Here, we used a combination of cosedimentation assays and microscope-based filamentous actin (F-actin) bundling assays to show that CLIP-170 can bind directly to F-actin *in vitro*. By studying CLIP-170 fragments and mutants, we found that the F-actin-binding domain is independent of the formin-activating FEED domain, requires the second CLIP-170 CAP-GLY motif (i.e., CAP-GLY 2) for efficient binding, and overlaps with the MT binding surface of the CAP-GLY domain. Consistent with these observations, our competition assays further indicate that CLIP-170 cannot bind to F-actin and MTs simultaneously. CLIP-170 overexpression did not have obvious effects on actin localization or morphology, initially arguing against the physiological significance of CLIP-170:F-actin interactions. However, effects of CLIP-170 overexpression might also be expected from the characterized indirect interactions between CLIP-170 and actin as described above, indicating that this observation is difficult to interpret. As a parallel approach to studying the potential significance of CLIP-170-actin interactions, we used bioinformatics and identified a residue in CAP-GLY 2 that is important for binding to F-actin but not for binding to MTs. This residue is well conserved across a range of organisms within CAP-GLY 2 but not between CAP-GLY 2 and CAP-GLY 1, consistent with the idea that the interaction between CAP-GLY2 and F-actin is functionally significant. Previously, we characterized direct binding of the core +TIP EB1 to F-actin and proposed that this binding may function to reduce EB1-binding to MTs and thus destabilize MTs in the actin-rich periphery of the cell (24). We speculate that binding of CLIP-170 to actin may function similarly, helping to destabilize MTs in at the cell periphery.

## Results

### The N-terminal “head” domain of CLIP-170 binds to F-actin directly

To understand whether CLIP-170 might have a more direct role in actin-MT crosstalk, our first approach was to test whether CLIP-170 binds directly to filamentous actin (F-actin) *in vitro* using high-speed cosedimentation assays. We chose the CLIP-170 N-terminal fragment H2 (Figure 1A; also known as the CLIP-170 MT-binding “head” domain fragment #2) for this test because the full-length CLIP-170 protein is easily degraded *in vitro* (25) and is auto-inhibited (26). After subtracting the background ~4% of H2 that self-pelleted in the absence of F-actin, we found that ~26% of H2 moved into the pellet in the presence of F-actin (3 μM) in our initial F-actin binding assays (Figure 1B). These results suggested that CLIP-170 can bind to F-actin directly via its N-terminal head domain.

**Figure 1.**
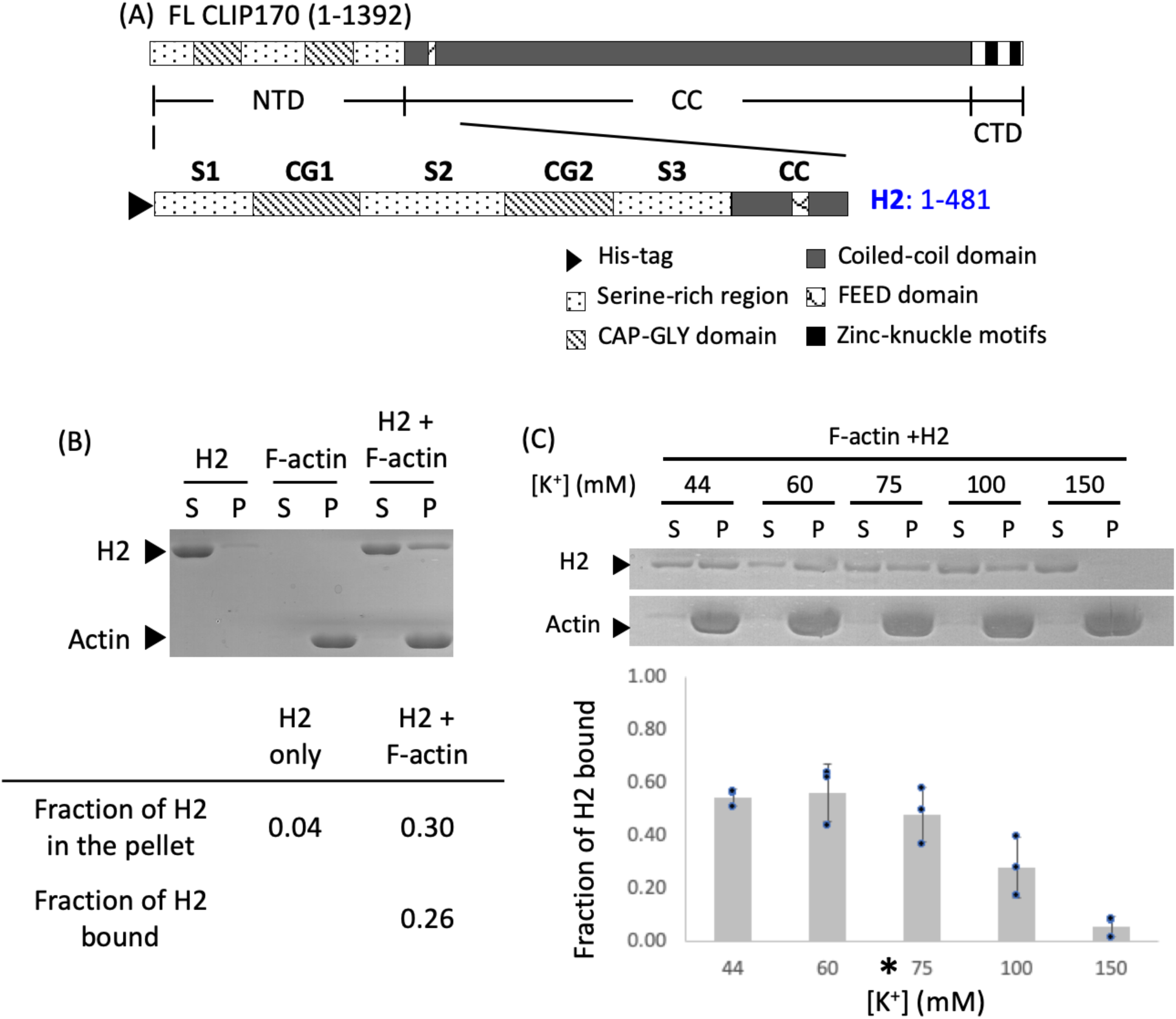
CLIP-170 binds to F-actin directly through its N-terminal head domain in an electrostatic-dependent manner. (A) Diagram of full-length CLIP-170 and the H2 fragment. (B) Binding of H2 to F-actin. High-speed cosedimentation assay with H2 (4 μM), phalloidin (0.8 μM), and pre-polymerized F-actin (3 μM) in PEM50 buffer; S indicates the supernatant; P indicates the pellet. The table shows an example of how the ‘fraction of H2 bound’ was calculated in this manuscript (see Methods). (C) Effect of changing salt concentration on cosedimentation of H2 (4 μM) with phalloidin-stabilized F-actin (6 μM). Salt concentration here corresponds to the total potassium concentration in the reaction (see Methods). * indicates the salt concentration of PEM50 pH 6.8, which is the standard buffer used in this manuscript. These data show that in the presence of increased salt, H2 in the pellet decreased whereas H2 in the supernatant increased. Error bars represent the standard deviation (n=3).

To better understand this CLIP-170:F-actin interaction, we performed a salt-sensitivity test. The fraction of H2 bound to 6 μM F-actin decreased with increasing salt concentration (Figure 1C). This observation indicates that ionic interactions play a role in the CLIP-170:F-actin interactions. For comparison, the physiological potassium ion concentration is ~150 mM (27). For the rest of the work in this manuscript, the experiments used a buffer (PEM50) that has a potassium ion concentration equivalent to 75 mM. We chose this buffer because it allowed us to work with moderate F-actin concentrations and is consistent with salt concentrations used for other F-actin binding studies in the literature (e.g., (28)).

Taken together, these results indicate that the N-terminal domain of CLIP-170 can bind F-actin directly, and that this interaction is mediated at least partly by ionic interactions.

### The F-actin-binding and MT-binding regions of CLIP-170 overlap with each other

To determine which region(s) of CLIP-170 are responsible for the CLIP-170:F-actin interaction, we measured the K_D_ of various CLIP-170 N-terminal constructs with F-actin using high-speed cosedimentation assays (Figure 2). These constructs were H2, as well as various CLIP-170 fragments containing one of the CG (CAP-GLY) domains, with or without their nearby S regions (serine-rich regions) (Figure 2A). H2 is a dimer, and other CLIP-170 fragments are monomers (25,29). Note that we did not include H1 (CLIP-170 N-terminal fragment with no coiled-coiled domain) in this test because the molecular weight of H1 is too similar to actin and so the two proteins cannot be resolved with SDS-PAGE.

**Figure 2.**
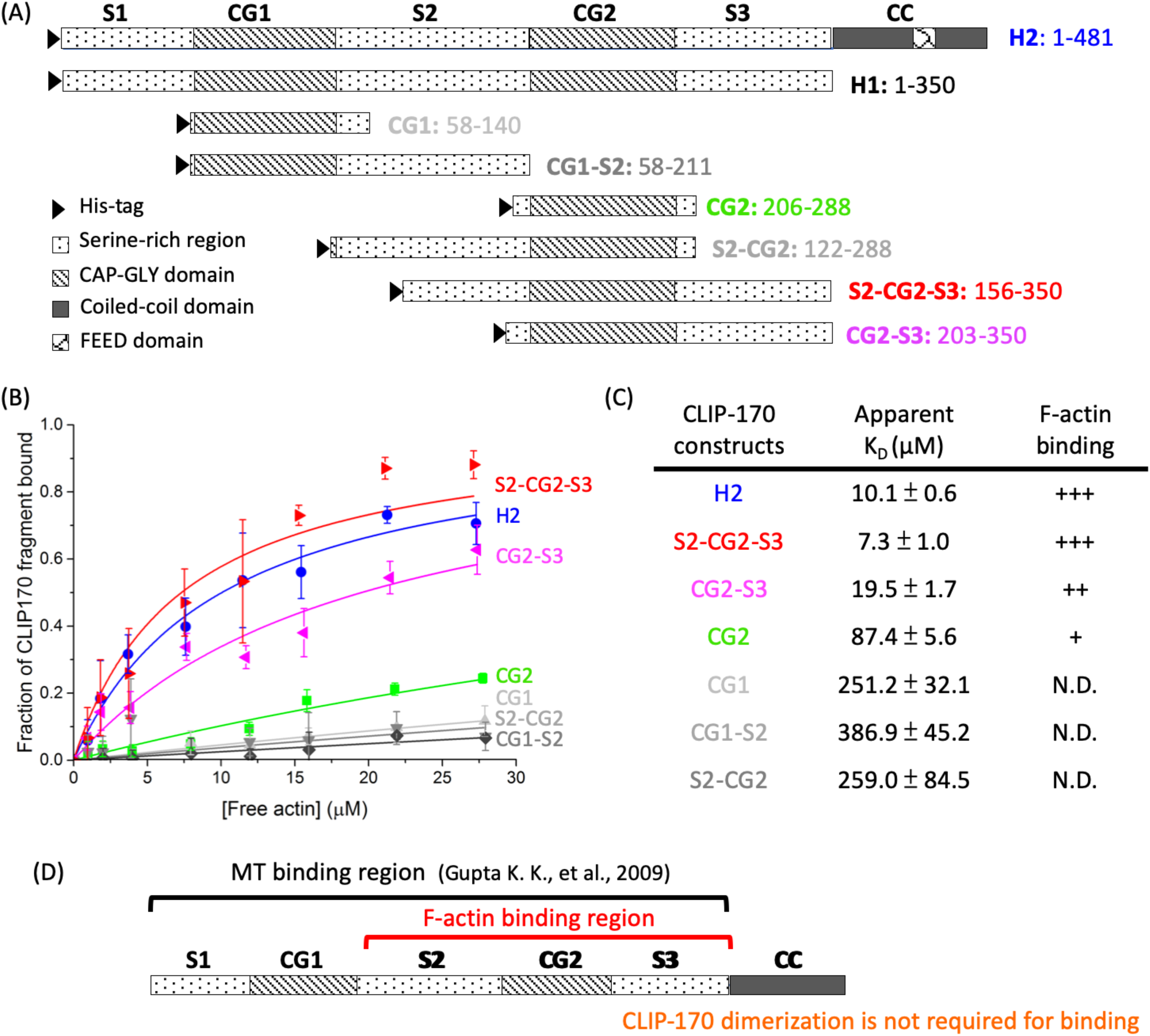
CLIP-170 binds to F-actin directly via CG2 and the surrounding serine-rich regions. (A) Summary of CLIP-170 fragments used in this study. (B, C) Determining the K_D_ of CLIP-170 fragments. High-speed cosedimentation assays with CLIP-170 fragments (4 μM), phalloidin (0.8 μM), and F-actin (concentrations as indicated) in PEM50 buffer generated the binding curves in (B). Curves were fitted to the data by OriginPro (simple binding equation with assumption of a 1:1 binding ratio). Error bars are standard deviation with n=3. The apparent K_D_ values of CLIP-170 fragments were extracted from the curve fits and are listed in the table (C). N.D., binding not detected. (D) Summary of the CLIP-170 regions involved in binding to F-actin and MTs. This diagram shows that the CLIP-170 F-actin-binding and MT-binding regions overlap.

No binding was observed between F-actin and constructs lacking CG2 (Figure 2 B, C). In contrast, all constructs containing CG2 bound to F-actin. Cosedimentation assays indicated that the K_D_ of F-actin:CG2 (~90 μM) was ~9-fold weaker than that of H2 (~10 μM) (Figure 2 B, C). With CG2-S3, the F-actin-binding affinity recovered to 20 μM (~2-fold weaker than that of H2), and having both nearby S regions (S2-CG2-S3) fully restored the F-actin binding affinity to a value similar to H2 (~10μM) (Figure 2 B, C).

These observations led to several conclusions: (1) the minimal F-actin-binding region for full-binding activity corresponds to CG2 and its bilateral serine-rich regions (S2-CG2-S3) (Figure 2D); (2) the F-actin-binding region does not include the FEED domain identified as binding to formins (23), so this interaction is different from that involved in formin-dependent actin polymerization; (3) dimerization is not required for this interaction because S2-CG2-S3 has no coiled-coil domain; (4) the two CG domains are not equivalent – the CG2 subdomain plays a more important role in mediating F-actin-binding than does the CG1 subdomain. In considering conclusion (4), it is interesting to note that CG2 is also more important for binding to MTs (30,31).

### Known MT-binding surfaces of CLIP-170 have positively charged basic grooves that are highly conserved

The observation that the CLIP-170 F-actin-binding region overlaps with the stronger of the two CLIP-170 MT-binding regions brings up the question of whether CLIP-170 residues directly involved in binding to MTs are also involved in binding to F-actin. To answer this question, we identified the residues previously established as binding to MTs (30) and highlighted them on the crystal structures of the two CG domains (CG1 (PDB:2E3I (30)) and CG2 (PDB:2E3H (30))). For each CG domain, we defined as the “front” the side that has the most residues identified as being involved in MT-binding (Figure 3A). In particular, note that the front side contains the well-conserved GKNDG sequence and an adjacent groove that binds to the EEY/F motif of tubulin; this surface also binds to the CCHC motif of the zinc-knuckle domains during CLIP-170 autoinhibition (30,32).

**Figure 3.**
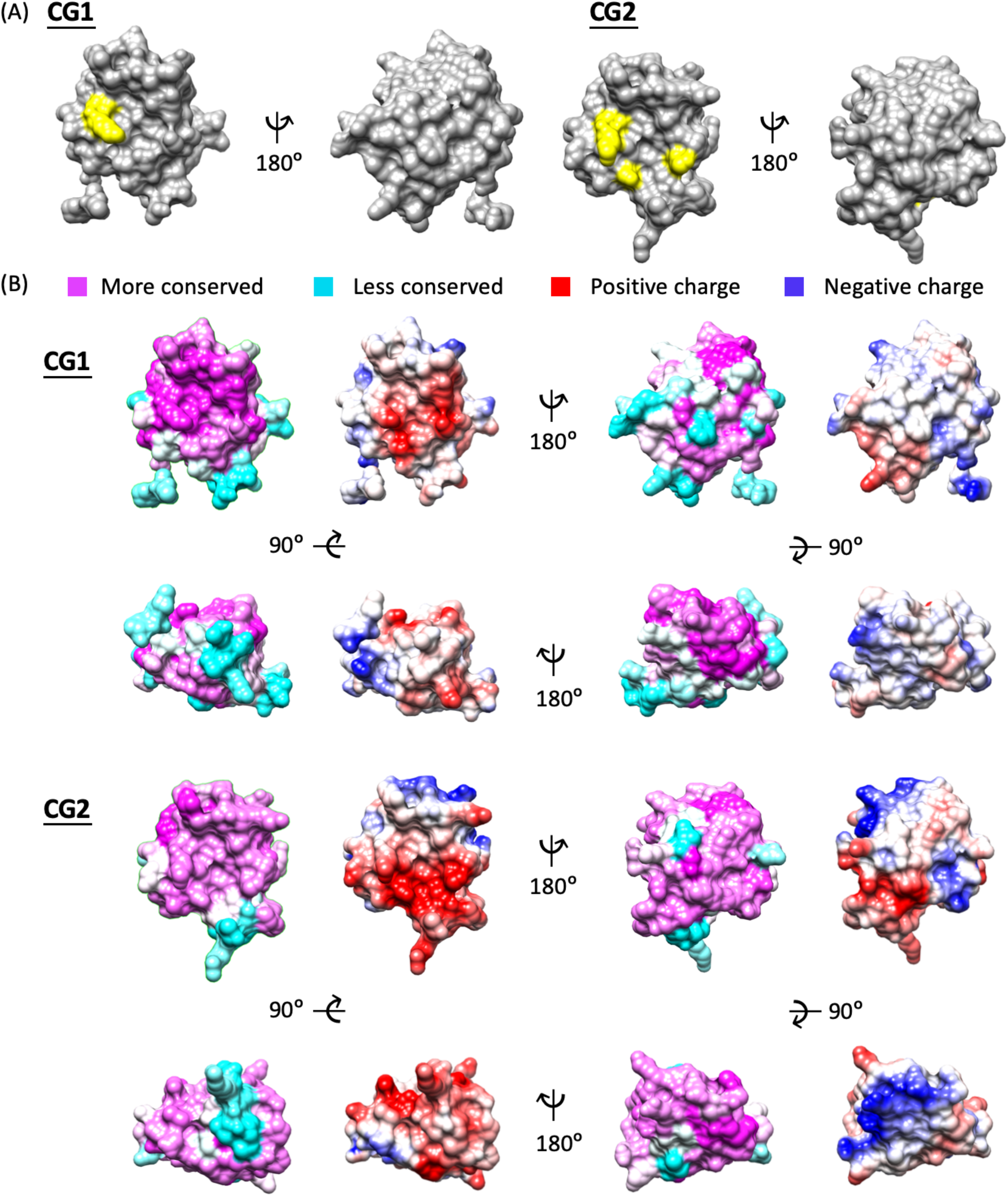
Conservation and electrostatic distribution of CLIP-170 CG1 and CG2 domains. (A) Diagram of known tubulin-binding sites. Tubulin-binding residues found in the literature (30) were plotted in yellow on the human CLIP-170 CG1 (PDB:2E3I) and CG2 (PDB:2E3H) crystal structures (30), which were aligned using Chimera (51) to display the same faces. (B) Conservation and electrostatic maps of CG1 and CG2. Sequence conservation across a range of vertebrates (see Methods) and electrostatic distribution in human CLIP-170 were mapped onto the CG1 and CG2 structures described in (A). Note that some amino acids in cyan (less conserved) at the bottom part of the CG domains are shown as poorly conserved due to alternative splicing in some organisms (e.g. fish) (see alignment in Supplementary Information).

To obtain more information about the F-actin binding surface of CG2, we used conservation mapping. We first gathered CLIP-170 sequences from a range of vertebrate organisms and then plotted the amino acid conservation as observed across these organisms onto each of the two human CG domains. In parallel, we also plotted the electrostatic distribution for the human CG domains (see Methods) (Figure 3B). We observed that for CG1, the front side is more conserved than the back side, while for CG2 both the front and back sides are highly conserved. On the front side of CG2, the basic EEY/F binding groove near the GKNDG motif previously mentioned is highly conserved, consistent with the evidence that this groove participates in tubulin binding. We noticed that when plotting the two structures by the electrostatics of the residues, the front sides of both CG domains are highly positively charged especially around the groove, while the back sides are generally neutral or negatively charged (Figure 3B). The observation of highly conserved and positively charged basic grooves on the front sides of both CG1 and CG2 agrees with previous studies (30).

The sum of this structural analysis and the results of our experiments led us to hypothesize that the residues in the positively charged groove of CG2 are involved in binding to F-actin. The key pieces of data leading to this hypothesis are that the F-actin binding region of CLIP-170 contains CG2 (but not CG1) (Figure 2), the CLIP-170:F-actin interaction partly depends on ionic interactions (Figure 1C), and F-actin is net negatively charged (33).

### Residues near the positively charged groove were selected for subsequent study by site-directed mutagenesis

To test whether CLIP-170 binds to F-actin through the front groove of CG2, we selected and mutated highly conserved and positively charged residues on this surface. In parallel, we mutated similarly positioned residues in CG1, expecting these mutants to serve as negative controls.

The residues of the CG2 groove have been well-studied for their MT-binding interactions (30). Early work demonstrated that the K224A, K252A, and K277A mutants of human CLIP-170 have reduced MT-binding ability, while K238A, K268A, R287A, and R298A do not interfere with MT-binding (30). We included 4 of these mutants in our study of F-actin-binding (K224A, K238A, K252A, and K277A), but we excluded K268A, R287A, and R298A because these three positions are further away from the basic groove. In addition, we selected a series of other positive residues located near the basic groove and mutated them to alanine. Those mutations are K70A, K123A, and H276A. Finally, we mutated the K and N residues of the highly conserved GKNDG motif of the CG domains. Previous work has shown that these two residues facilitate MT tracking (15,34) and EEY/F binding (30,32). Thus, we also included mutants of N98E, K98E-N99D, K252A, N253A, and K252E-N253D in our study to test their role in F-actin-binding.

We chose to make the mutations in the H2 fragment of CLIP-170 because the molecular weight of the H1 fragment is too similar to actin to allow separation from actin in SDS-PAGE. This strategy works for testing the effects of the mutations in F-actin binding assays; however, it leads to problems in determining the effects of mutation on MT-binding because the strong affinity and multivalent binding between H2 and MTs makes it difficult to measure the affinity differences via cosedimentation assays. Thus, we adapted a strategy previously used to identify residues involved in MT-binding (e.g., (30)). Specifically, we made the same mutations in CG1 or CG2 fragments (which have weaker CLIP-170:MT affinity) and quantitatively assessed the binding of these fragments to MTs. In total, at the end of this mutation-design process, we had 9 single mutants and 2 double mutants in H2 and CG fragments for the subsequent F-actin or MT-binding assays, respectively (Figure. 4 A, B, and Figure S1).

**Figure 4.**
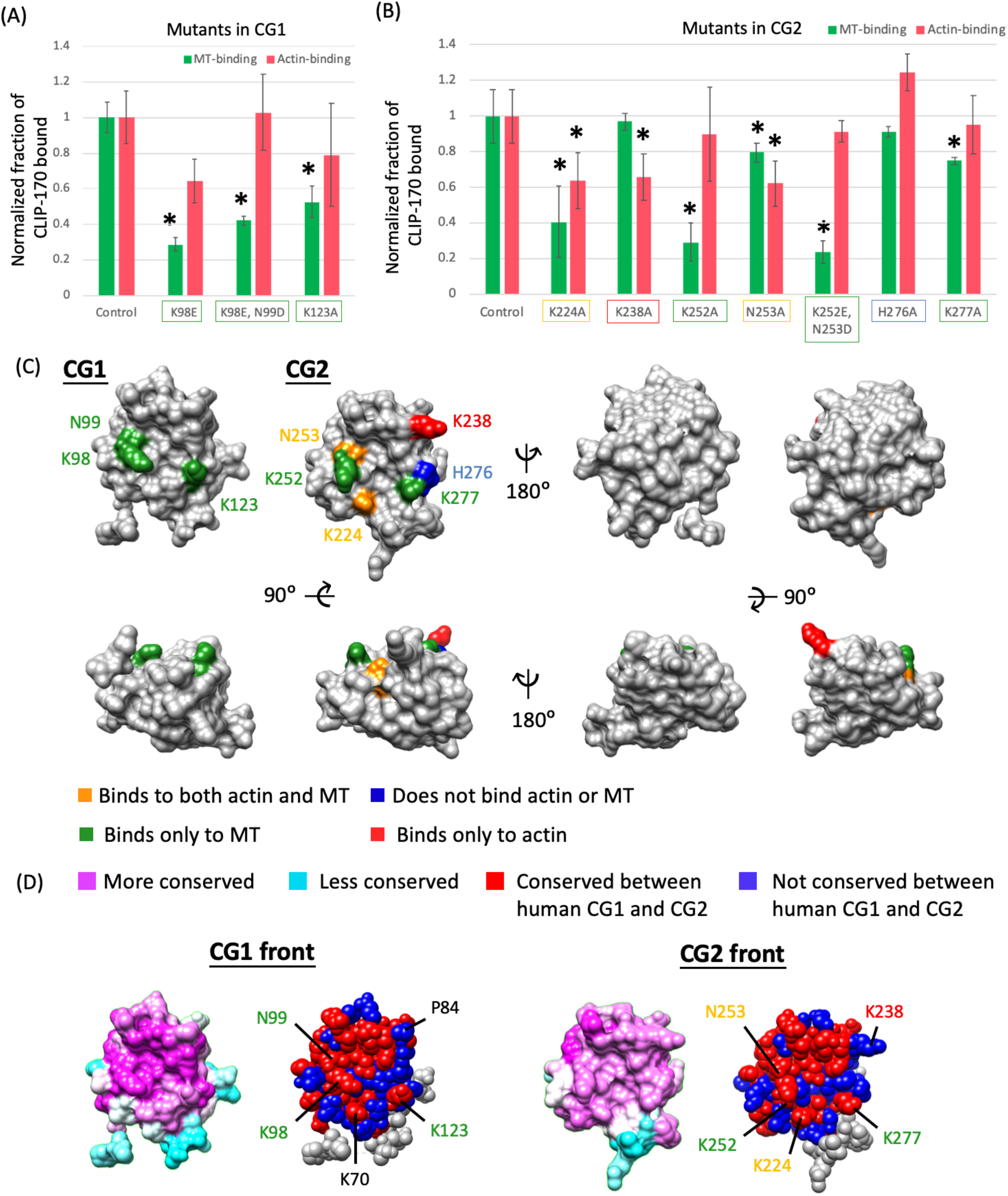
Characterization of CLIP-170 and CLIP-170 mutants. (A, B) Summary of F-actin and MT-binding activity of selected CLIP-170 mutants as determined from F-actin or MT-binding assays. For F-actin binding assays, 2.3 μM H2 or H2 mutants and 10.5 μM F-actin were used. For MT-binding assays, 5 μM CG1, CG2, or their mutants and 10.5 μM Taxol-MTs were used. For ease of comparison, the fraction of protein bound (Figure S1) was normalized against the corresponding control (H2, CG1, or CG2) to represent the binding activity (see Methods). Error bars represent standard deviation. For each mutant, n=3-4. For controls, n= 6 (for CG1), 9 (CG2), and 22 (H2). * indicates p-value < 0.05; as described in the results, only the subset of controls corresponding to that experiment was used to determine the indicated p-values. (C) Summary of the relative positions of residues involved in F-actin- and MT-binding as highlighted on the CG1 and CG2 structures. (D) Summary of the amino acid differences between human CG1 and CG2. The magenta-cyan conservation maps on the left summarize conservation within each CG domain and were reproduced from Figure 3. The red-blue maps on the right indicate amino acids which are identical (red) or different (blue) in the alignment of the two human CG domains. Amino acids in gray are outside of the alignment regions. Text colors indicate amino acids that contribute to MT binding (green), F-actin binding (red), and binding to both filaments (orange); black text indicates a residue in CG1 that was not analyzed because it has a serious self-pelleting problem, though the analogous position in CG2 is involved in binding to both MTs and actin.

### Analysis of CG mutants suggests that the F-actin and MT-binding surfaces overlap with each other

To test whether these selected residues in the CG1 and CG2 grooves are important for binding to F-actin and/or MTs, we performed high-speed cosedimentation assays with our CLIP-170 mutants. Wild-type H2, CG1, or CG2 were used as controls. Because high-speed cosedimentation assays can be prone to technical problems (e.g., self-sedimentation or loss of protein from sticking to tube walls), we always ran both positive and negative controls in parallel to experimental tests to ensure that experiments run at different times could be compared. However, it is important to note that this strategy results in a much larger sample number for control groups than the mutant groups, which can artificially reduce the p-value when comparing the affinity of mutant proteins with the wild-type proteins (35,36).

To avoid this problem of artificially reduced p-values, we included only the control that was run in parallel with a particular mutant to calculate the p-value for that mutant. With this approach, three mutants had significantly reduced F-actin binding (K224A, K238A and K253A), and none had increased F-actin binding (Figure 4A, B and figure S1). For MT-binding assays, 8 mutants had significantly decreased MT-binding affinity (Figure 4 A, B and Figure S1).

Before interpreting these data in detail, all mutants were tested by circular dichroism (CD) for secondary structure to assess whether they were properly folded. As shown in Figure S2, the shapes of the CD spectra for all mutants were very similar to those of the wild-type controls. Because of this similarity, no mutants other than K70A (which showed a serious self-pelleting problem) were excluded from further analysis. However, we do note that the magnitude of the CD spectrum was somewhat different for a few mutants (H2-K70A, H2-K224A, and potentially the CG1-K98E,N99D and CG2-N252A mutants), which may indicate some level of misfolding (Figure S2).

Consideration of all these data together leads to the following conclusions: of the 11 mutants tested, 2 mutants have significantly reduced binding to both F-actin and MTs (K224A and K253A); 6 mutants have significantly reduced binding only to MTs (K98E, K98E-N99D, K123A, K252A, K252E-N253D, and K277A); 1 mutant has significantly reduced binding only to F-actin (K238A); and 1 mutant has no significant change in binding to either F-actin or MTs (H276A) (Figure 4 and S1). As expected, all of the residues involved in binding to F-actin are in CG2. The observation that residues involved in binding F-actin and MTs are on the same surface of CLIP-170 CG2 indicates that the F-actin and MT-binding surfaces of this CG domain overlap with each other.

### Conservation patterns support a role for residue K238 in binding to F-actin

The sequences of human CG1 and CG2 are 59% identical. It is well-established that both CG1 and CG2 bind to MTs, while the data discussed above indicate that binding to F-actin is mediated primarily by CG2 (Figures 2–4). These data led us to hypothesize that amino acids involved in MT binding would be conserved between CG1 and CG2, but those involved in F-actin binding would be different between the two CG domains. To test this idea, we aligned the two human CG1 and CG2 sequences and mapped the alignment result on the CG1 and CG2 protein structures to show amino acids that are identical (red) and different (blue) between the two CG domains (Figure 4D). The results agree with our predictions that the MT-binding residues are conserved (indeed, identical) between the two CG domains, while the F-actin-binding residue (K238) is different from its corresponding position in CG1 (P84). Interestingly, both K238 and P84 are conserved across vertebrates (Figure S3A), which suggests that they are both functionally significant.

In summary, we found that our analysis of MT-binding residues agrees with previously published work (30). We extended the understanding of those residues by testing the ability of mutants in these residues to bind F-actin. In addition, we also evaluated mutants that were not included in the previous literature. The sum of these data indicates that the CG2 of CLIP-170 is important for F-actin binding, and that the F-actin binding surface of the CLIP-170 CG2 domain overlaps with its MT-binding surface.

### CLIP-170 bundles F-actin *in vitro*, and bundling activity correlates strongly with F-actin binding

Because the F-actin and MT-binding surfaces of CLIP-170 appear to overlap (Figure 4), we hypothesized that MTs might compete with F-actin for binding to CLIP-170. Unfortunately, high-speed cosedimentation assays are not suitable for testing this hypothesis because both MTs and F-actin will be pelleted down, and thus it would not be possible to determine whether CLIP-170 is bound to actin, MTs, or both. Previous studies have demonstrated that the formation of F-actin bundles can be used as a read-out for protein binding (24). We decided to try to use a similar strategy to evaluate the possible competition between F-actin and MTs for binding to CLIP-170.

CLIP-170 has been shown to induce MT bundles both *in vitro* (37) and *in vivo* (38); however, it was unknown whether CLIP-170 can induce F-actin bundles. Thus, we first incubated H2 and fluorescently labeled F-actin together to evaluate the F-actin bundling activity of CLIP-170 *in vitro*. The results indicated that CLIP-170 can induce the formation of F-actin bundles (Figure 5).

**Figure 5.**
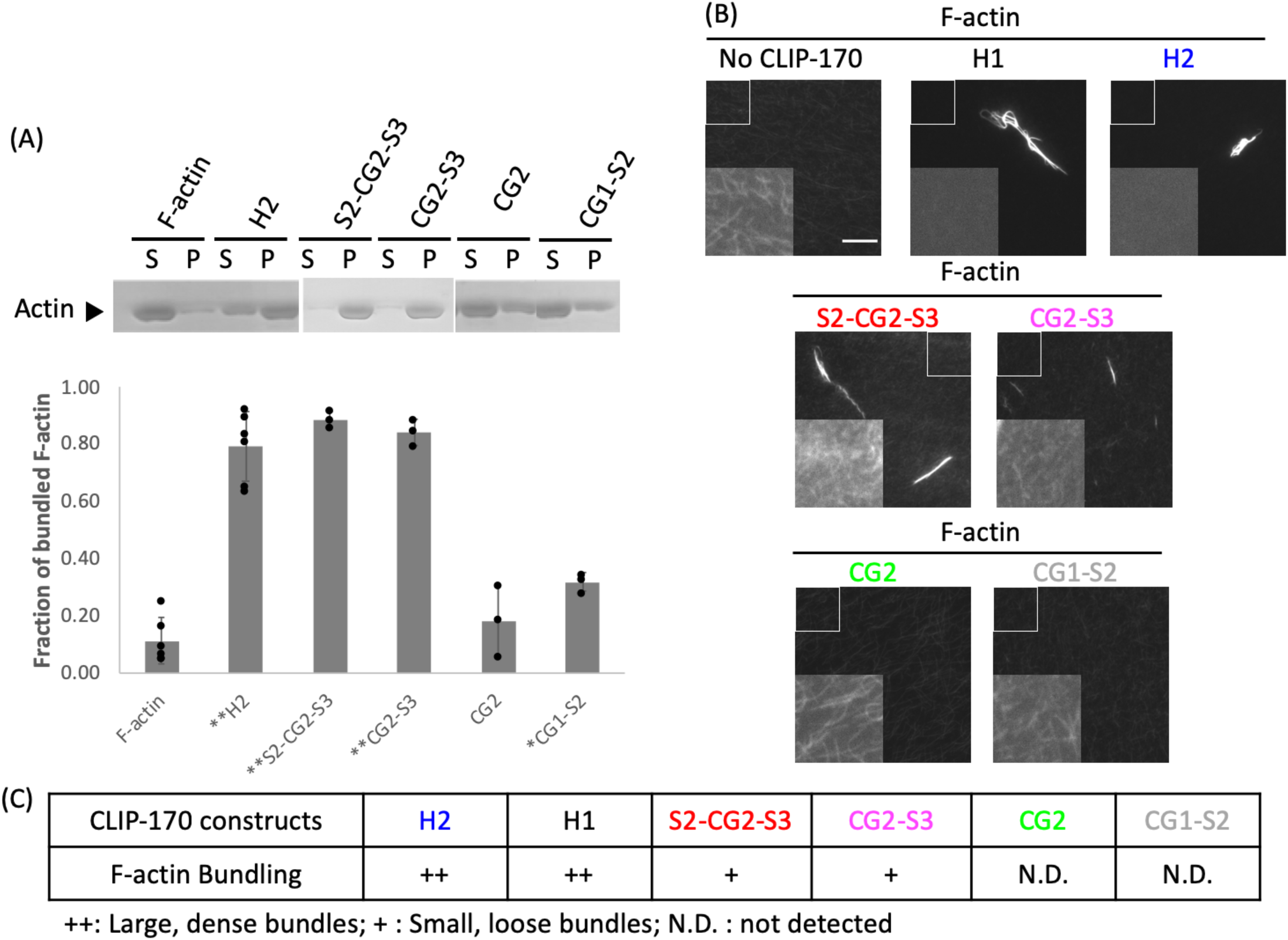
CLIP-170 triggers formation of F-actin bundles in vitro. (A) Low-speed cosedimentation assay with 4 μM CLIP-170 fragments, 5 μM F-actin, and 0.8 μM Alexa-488-phalloidin in PEM50, performed to assess the F-actin bundling ability of CLIP-170 fragments. S indicates the supernatant that contains F-actin; P indicates the pellet that contains F-actin bundles. (B) CLIP-170 fragments induced different F-actin bundle phenotypes. The fluorescence microscope images show that H1 and H2 induced large dense F-actin bundles; S2-CG2-S3 and CG2-S3 induced small loose bundles. CG1-S2 and CG2 had no or little ability to bundle F-actin. The main images were normalized to a common level chosen to best visualize bundles; the insets were normalized to a common level chosen to best visualize individual filaments. Scale bar = 10 μm. (C) Table summarizing the F-actin bundling ability of CLIP-170 fragments based on these data.

Next, we tested whether the F-actin bundling activity and the F-actin binding activity are functionally separable by incubating CLIP-170 fragments that bind to F-actin (H2, H1, S2-CG2-S3, CG2-S3, and CG2) (Figure 2) with fluorescently labeled F-actin. In parallel to this visual assay (Figure 5B), we performed more quantifiable low-speed cosedimentation assays, under the assumption that bundled F-actin will go into the pellet, while unbundled F-actin will stay in the supernatant (Figure 5A). As predicted, only CLIP-170 fragments that bind well to F-actin can bundle F-actin, as assessed by either assay (Figure 5). This observation suggested that F-actin bundling activity can be used as a read-out for CLIP-170:F-actin binding.

### MTs and tubulin dimers appear to compete with F-actin for binding to CLIP-170

We used these CLIP-170 F-actin bundling/binding activities to investigate whether MTs compete with F-actin for binding with CLIP-170. Briefly, we developed an assay in which we incubated the CLIP-170 fragments with MTs or tubulin dimers first, then added Alexa-488 phalloidin-labeled F-actin. The assumption of this assay is that if MTs compete with F-actin for binding to CLIP-170, the ability of CLIP-170 to bundle F-actin should be reduced in the presence of MTs.

Our results showed that CLIP-170 fragments (H1 and H2) bundle F-actin, as expected from our earlier experiments (Figure 5), and that the F-actin bundles disappeared in the presence of high (10 μM) concentrations of either MTs or tubulin dimers (Figure 6 A,B). These results implied that MTs and F-actin compete for binding to CLIP-170, but to follow up on this initial conclusion, we did a more in-depth experiment by adding different amounts of MTs or tubulin in the competition assays. We observed that the F-actin bundles decreased in size and eventually disappeared as the concentration of MTs or tubulin increased from 0-5 μM (Figure 6C).

**Figure 6.**
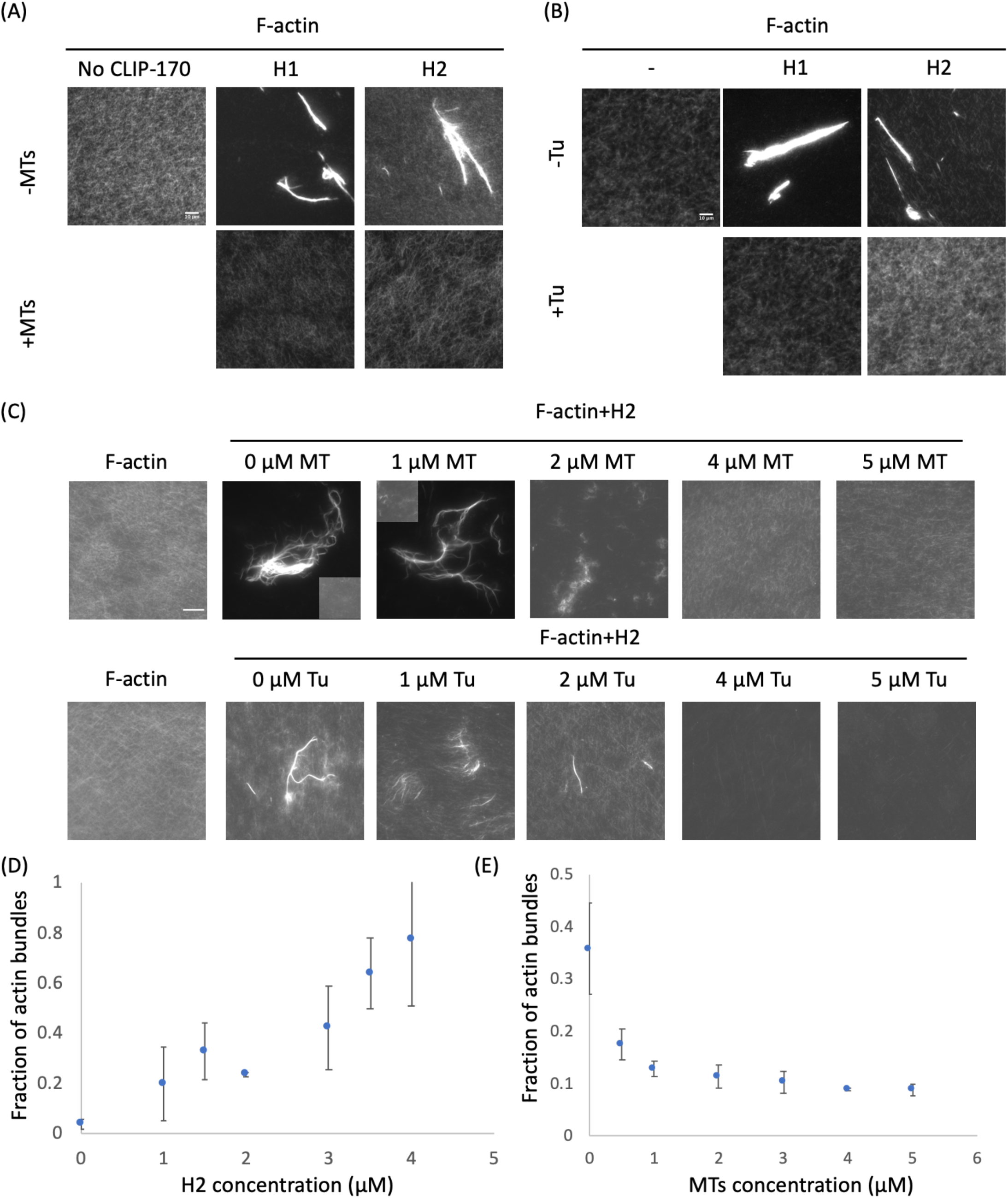
F-actin and MTs/tubulin compete for binding to CLIP-170. (A,B) Preincubation with high concentrations of MTs or tubulin (Tu) prevents interactions between F-actin and CLIP-170. To generate initial CLIP-170:MTs/tubulin interactions, 10 μM MTs or tubulin (as indicated) and 2 μM CLIP-170 fragments were mixed in PEM50. Alexa-488 phalloidin-labeled F-actin (2 μM) was then added to the premixed CLIP-170:MT solution 10 mins before imaging. Images in the same panel were adjusted to the same levels. (C) Competition assays were prepared as described in but with varying amounts of MTs or tubulin (Tu). Images in the same row were adjusted to the same levels except the second and third images from the left of row 1. These two images were adjusted to allow the visualization of bundles, and the inset is normalized the same as the other images to show unbundled F-actin in the background. (D) Low-speed cosedimentation assays with 2 μM F-actin and varying concentrations of H2 to generate a dose-dependent F-actin bundling curve. (E) Competition reactions were prepared as described in (A) but with various concentrations of MTs, 2 μM H2, and 2 μM F-actin. In order to quantify the competition assays, the reactions were analyzed by low-speed cosedimentation assays rather than microscopy. Error bars are standard deviation (n=3).

In order to obtain more quantitative data, we investigated the possible competition between F-actin and MTs by performing low-speed cosedimentation assays. As discussed above, these assays assume that bundled F-actin will sediment when centrifuged at low speed, but unbundled F-actin will not. First, we conducted experiments to determine whether the amount of F-actin bundling by CLIP-170 depends on the concentration of CLIP-170. As expected, our results show that F-actin bundles increase with the amount of H2 added (Figure 6D). Next, we tested the impact of pre-incubating the CLIP-170 with MTs. We observed that there was a dramatic inverse relationship between the concentration of MTs used in the assay and the amount of bundled F-actin (Figure 6E). Taken together, these results indicate that both tubulin and MTs compete with F-actin to bind to CLIP-170.

In conclusion, the results of our experiments show the CLIP-170 head domain can bind F-actin directly. This CLIP-170:F-actin interaction is mediated by the CG2 domain and its nearby S regions. Our data also indicate that the CLIP-170 F-actin-binding and MT-binding surfaces overlap; consistent with this idea, F-actin and MTs compete for binding to CLIP-170, meaning that the CLIP-170 head domain cannot bind to both filament types simultaneously.

### Overexpression of full-length CLIP-170 has no obvious effects on the actin cytoskeleton *in vivo*

To test whether the CLIP-170:F-actin interaction can be detected in cells, we overexpressed full-length GFP-CLIP-170 in COS-7 cells and stained for F-actin. We were interested to see if the proteins colocalized, and also whether CLIP-170 overexpression altered F-actin concentration or morphology. We observed that there are some areas of possible colocalization between CLIP-170 and F-actin in cells expressing low-levels of CLIP-170 (Figure 7). It is interesting to note that previous work has reported that CLIP-170 and actin colocalize at sites of phagocytosis (22). However, we observed no obvious colocalization between F-actin and CLIP-170 in cells expressing CLIP-170 at medium and high levels of overexpression (Figure 7). In addition, we did not observe obvious differences in actin morphology between untransfected cells and those overexpressing CLIP-170 (Figure 7).

**Figure 7.**
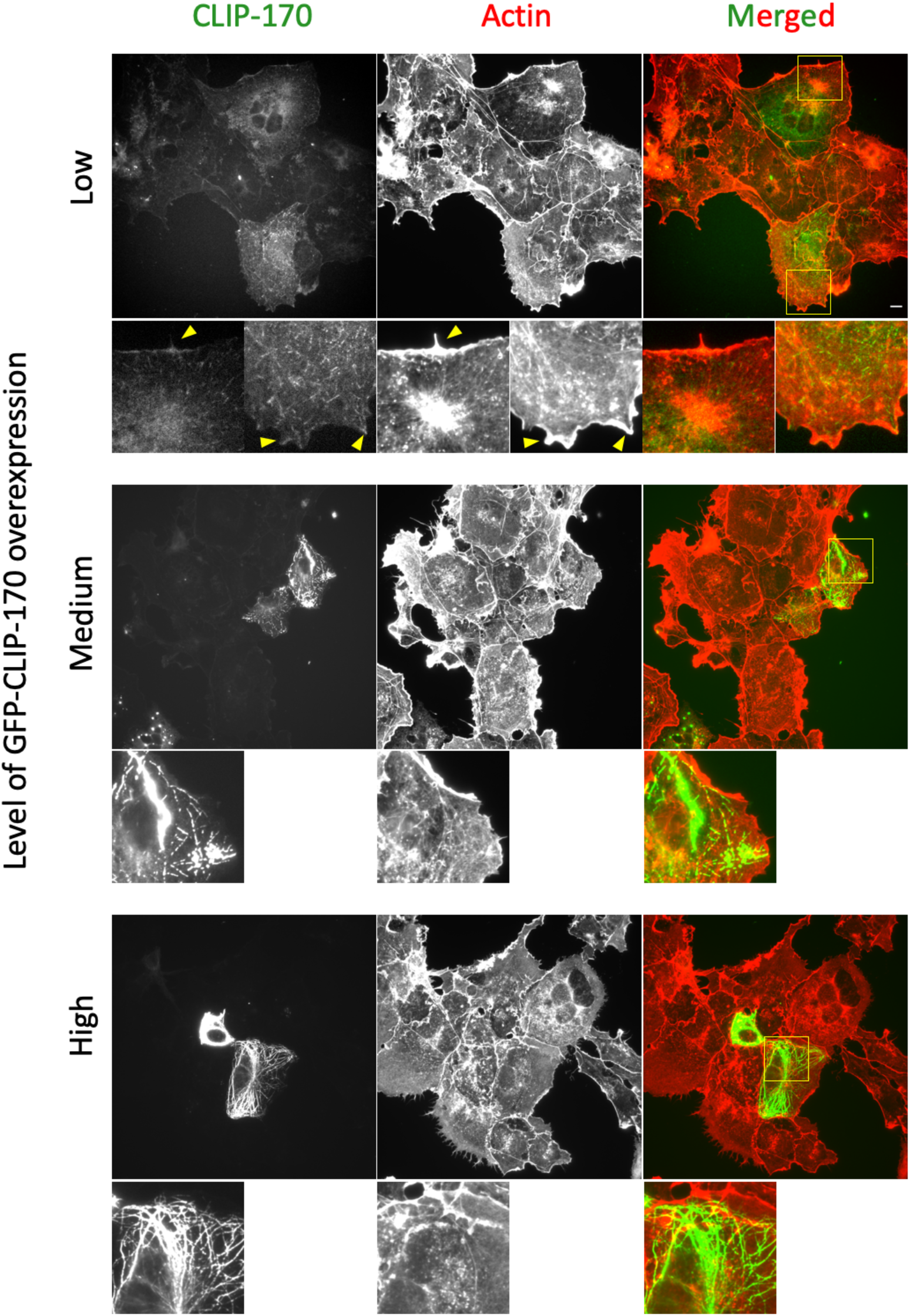
Overexpression of full-length CLIP-170 in cells has no obvious effect on the actin cytoskeleton. COS-7 cells were transfected to overexpress GFP-labeled full-length CLIP-170. Cells were fixed with PFA, and F-actin was labeled with rhodamine-phalloidin. F-actin and CLP-170 staining are shown at varying levels of CLIP-170 overexpression. Different levels of CLIP-170 overexpression lead to distinctive phenotypes in the MT cytoskeleton, including MT plus-tip labeling (low CLIP-170 expression), MT bundling and patch formation (medium expression), extreme MT bundling (high expression)(see also (39)). Smaller images shown in the bottom of each row are the zoom-in images of the yellow box(s). Note that there are some areas of colocalization visible in cells expressing low levels of CLIP-170 (yellow arrows). However, while there might appear to be some differences in F-actin staining between CLIP-170-transfected cells and nearby cells, we found that we could not reliably identify CLIP-170-transfected cells by looking only at the actin channel (representative images shown here), indicating that CLIP-170 overexpression does not cause obvious effects on actin morphology.

At first glance, these observations argue against a physiological role for the CLIP-170:F-actin interactions. However, the lack of obvious effects of CLIP-170 overexpression on the actin cytoskeleton is surprising because published evidence indicates that CLIP-170 activates formins (22,23), and because CLIP-170 is expected to impact actin through IQGAP (20,21). One possible explanation for the apparent lack of effect is that actin assembly is highly regulated, and this regulation is able to overcome any perturbation (direct or indirect) caused by CLIP-170 overexpression.

Because it would be very challenging to separate any direct effects of CLIP-170:F-actin interactions from indirect effects mediated by CLIP-170:formin or CLIP-170:IQGAP interactions, we decided to further investigate the question of physiological significance through studies of CLIP-170 conservation.

### CLIP-170 F-actin binding residues are highly conserved

If CLIP-170:F-actin interactions are functionally significant, one would expect that CLIP-170 F-actin-binding residues would be well-conserved throughout diverse species. To investigate this question, we selected 7 organisms from human to yeast, and we aligned their CG domains (Figure S2A). We observed that almost all MT-binding residues (K98E, N99D, K123, K224, K252, N253 and K277) are highly conserved in both CG1 and CG2 across species, as expected. The two residues (K224 and N253) that bind to both MT and F-actin are also conserved across species in both CG1 and CG2. The one residue that binds only to F-actin (K238) is also well-conserved as lysine in the CG2 of the animal CLIP-170 proteins, but the corresponding residue in CG1 is conserved as proline. This observation is consistent with our finding that CG2 binds to F-actin but CG1 does not. With regard to the fungal proteins, it is notable that the single CAP-GLY of the *S. pombe* Tip1p protein has lysine at this position, consistent with the idea that it too can bind to F-actin, but *S. cerevisiae* Bik1p has an alanine at this position, raising the possibility that Bik1 behaves differently (Figure S2A).

Overall, the observation of conservation in a CG2 residue (K238) that is involved in binding to F-actin but not MTs is consistent with the idea that F-actin binding is physiologically significant. We cannot exclude the possibility that this residue is conserved because of binding to other ligands, but the existing crystal structures argue against a role for K238 in binding to known CAP-GLY ligands (e.g. the CLIP-170 zinc knuckles (32,39), the C-terminal domain of SLAIN2 (40)), or EB1 (30).

We were interested to see that the position corresponding to K238 is conserved in both CG1 and CG2, but as different amino acids. We were curious to see what amino acid appears at this position in other CG-containing proteins, and whether this might provide insight into possible F-actin binding ability. To address this question, we aligned the CG domains of well-known CAP-Gly containing proteins from humans (Figure S3B). We speculated that if the amino acid position that corresponds to K238 is also a lysine, that protein might also have the ability to bind F-actin. CLIP-115 does have a lysine at this position, leading us to suggest that it may too bind F-actin. However, only one of the other proteins examined (CAP-350) has a positively charged amino acid at this position, leading us to speculate that few if any of the other CG domains bind to F-actin.

## Discussion

Our results show that the N-terminal domain of CLIP-170, which binds to MTs, can also bind F-actin directly (Figures 1 and 2). In addition, we found the binding surfaces of CLIP-170 to MTs and F-actin partially overlap with each other (Figures 4 and S1). Consistent with these observations, F-actin filaments were found to compete with MTs for binding to CLIP-170 in our bunding-based competition assays (Figure 6). We stress that while this actin bundling activity was useful as an experimental read-out, we are not suggesting that the bunding activity is physiologically relevant. It is interesting to note that dimerization of CLIP-170 is not required for the F-actin bundling activity (Figure 5). This observation may imply that there is more than one F-actin binding site in each of these constructs, but the observation that small peptides have been seen to bundle MTs (41) provides an argument against this interpretation. Although there is no obvious colocalization between CLIP-170 and F-actin in tissue culture cells (Figure 7), our bioinformatics studies show that both the MT and F-actin binding residues are well-conserved (Figure S3), consistent with the idea that binding of CLIP-170 to F-actin is functionally significant.

The K_D_ of this CLIP-170:F-actin interaction is ~10.5 μM when the salt concentration is half of the physiological level. This interaction is weak and may appear physiologically irrelevant. However, the concentration of actin at the cell cortex is extremely high (>300 μM) (42,43), which makes weak F-actin affinities potentially quite significant in the cortex regions. More specifically, if we assume that CLIP-170:F-actin binding is a simple interaction with K_D_ = ~20μM, and that the concentration of actin at the cortex region is 300 μM, more than 90% of CLIP-170 would be bound to F-actin in this environment. Thus, we suggest that the CLIP-170:F-actin interaction may be physiologically relevant in actin-rich regions such as the leading edge of migrating cells, where the actin concentration (>300 μM) (42,43) is much higher than the tubulin concentration (~20 μM in the cytosol as a whole) (44).

As discussed in the introduction, previous studies have shown that CLIP-170 regulates the actin cytoskeleton by binding to known actin-binding proteins such as IQGAP1 or formin (Figure 8) (20–23). Moreover, it was shown that CLIP-170 forms a complex with formin through the CLIP-170 FEED domain to promote actin polymerization (22,23). Our results show that the S2-CG2-S3 fragment, which does not contain the FEED domain, has binding activity similar to the H2:F-actin interaction (Figure 2). These results indicate that the CLIP-170:F-actin direct binding we found is independent of the FEED domain and raise the possibility that CLIP-170 can regulate the actin cytoskeleton through these direct interactions. While we cannot rule out this possibility, the observation that CLIP-170 overexpression does not have an obvious effect on actin morphology argues against this idea (Figure 7). Instead, we suggest that interactions between actin and CLIP-170 serve to downregulate the MT-promoting activity of CLIP-170 in actin-rich regions of the cell. Indeed, it is interesting to consider the possibility that promotion of actin polymerization by CLIP-170 helps to recruit CLIP-170 off of MT tips. Whether evidence can be found for such a model will be an interesting topic for future work.

**Figure 8.**
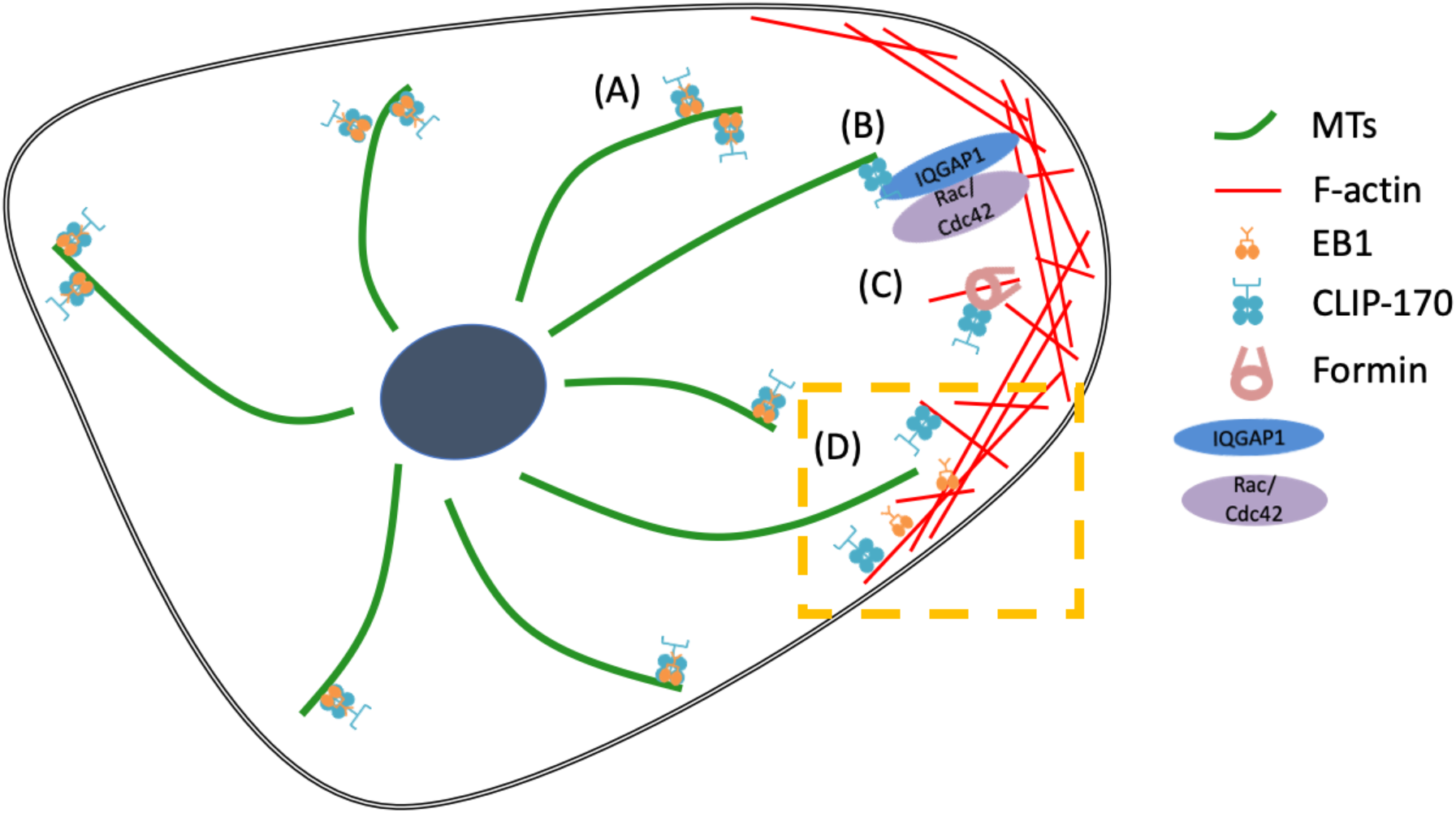
Current and proposed models of the role of CLIP-170 in cytoskeletal regulation. (A) EB1 and CLIP-170 track the MT plus end to regulate MT dynamics (53). (A) IQGAP1 connects the actin and MT cytoskeletons by associating with CLIP-170 (20,21). (C) CLIP-170 promotes actin polymerization through formins (22,23). (D) We propose that in the actin-rich cell periphery, the high concentration of actin competes CLIP-170 off of microtubules, promoting MT depolymerization in that region, similar to what was previously proposed for EB1 (24).

### Conclusions

In this manuscript, we showed that the microtubule plus-end tracking protein CLIP-170 can bind directly to actin via its second CAP-GLY motif, that this interaction is weak but strong enough to be relevant in the actin-rich cortex, and that binding to actin and MTs is mutually exclusive. Previously, our lab observed that another +TIP, EB1, can bind directly to MT and F-actin, and there is a competition between MT and F-actin filaments for EB1 (24). Based on these observations, we proposed that binding of EB1 to F-actin may cause EB1 to relocate from MTs to F-actin in the actin-rich cell cortex, thus promoting the destabilization of MTs near the cell edge (Figure 8) (24). Similarly, we have shown here that the +TIP CLIP-170 can bind directly MT to F-actin, and that MTs compete with F-actin for binding to CLIP-170. These observations lead us to suggest that CLIP-170:F-actin interactions may also function to destabilize MTs in the actin-rich cortex. Overall, we suggest that +TIP:F-actin binding may constitute a type of actin-MT crosstalk that destabilizes the MT cytoskeleton near the cell edge.

## Materials and Methods

### CLIP-170 constructs and protein purification

pET-15b-His-tagged CLIP-170 fragments used in this paper were described previously: H2 (H2^1–481^, (45)), H1 (H1^1–350^, (45)), CG1 (H1^58–140^, (29)), CG1-S2 (H1^58–211^, (31)), CG2 (H1^206-288^, (29)), S2-CG2 (H1^122-288^, (29)), S2-CG2-S3 (H1^156-350^, (31)), and CG2-S3 (H1^203-350^, (31)).

CLIP-170 site-directed mutants were generated in H2, CG1, or CG2 fragments depending on experiment needs. Residues were selected based on their conservation, electrostatics, and location information. Selected residues were mutated to alanine or oppositely-charged amino acids by PfuUltra II Hotstart PCR Master Mix (Agilent). All mutated sequences were confirmed by Sanger sequencing. His-tagged CLIP-170 fragments and mutants were expressed in BL21 (DE3) and purified by the standard His-tagged purification protocol from Novagen (69670-5, Sigma-Aldrich) with the following modifications. Briefly, cells were induced by isopropyl β-D-1-thiogalactopyranoside for 2 hr at 37°C and were harvested by centrifugation at 4000 × g for 10 min. Cell pellets were resuspended, sonicated, and centrifuged at 27,000 × g for 1 hr at 4°C before purifying with Ni2+ affinity chromatography. Eluted proteins were dialyzed in PEM buffer (100 mM PIPES, 2 mM MgCl, 1 mMEGTA, pH 6.8) with the reducing reagent β-mercaptoethanol (7 μL for 50 mL PEM buffer). Protein concentrations were determined by Bradford assays, and protein purity was assessed by separating samples on a 10% SDS-PAGE with a subsequent Coomassie stain. The concentrations of all CLIP-170 fragments were calculated as monomers, even though H2 forms dimers. All purified proteins had the expected molecular weight and purity in the purified solution fraction. Purified CLIP-170 constructs were stored at −80°C and thawed on ice before use.

### Actin and tubulin purification and polymerization

Tubulin was purified by two cycles of polymerization and depolymerization from porcine brain as described previously (45). Taxol-stabilized MTs were polymerized by the stepwise addition of Taxol (45). Both tubulin and MTs were stored at −80°C. MTs were thawed rapidly at 37°C immediately before use.

Globular actin (G-actin) was purified from rabbit muscle acetone powder (Pel-Freez Biological) by a cycle of polymerization and depolymerization as described previously (46). Purified G-actin was stored in a dialysis bag in calcium buffer G (2mM Tris-HCl, 0.2mM ATP, 0.5mM DTT, 0.1mM CaCl2, 1mM sodium azide, pH 8) and the buffer was refreshed weekly. To polymerize filamentous-actin (F-actin), G-actin was first converted to MG-actin with ME buffer (5 mM MgCl_2_ and 0.2 mM EGTA) for 5 min at room temperature. Then, KMEI buffer (50 mM KCl, 1 mM MgCl_2_, 1 mM EGTA, and 10 mM Imidazole-HCl (pH 7)) was added for 1 hr at room temperature to polymerize F-actin. The same process was performed with calcium buffer G to generate a complementary buffer for reaction without any F-actin as a negative control (Figure 1B).

### High-speed cosedimentation assays (binding assays)

The binding affinities of CLIP-170 fragments for F-actin or MT were assessed by high-speed cosedimentation assays. Briefly, CLIP-170 constructs, the relevant filament, and filament stabilizer (0.8 μM phalloidin for F-actin, or 10 μM Taxol for MTs) were mixed in buffers as described below (concentrations as indicated in the figure legends) and were incubated for 25 min followed by 15 min centrifugation at 184,000 × g. The temperature for incubation and centrifugation depended on which cytoskeletal filament was used. F-actin binding assays were performed at room temperature (~25 °C), and MT assays were performed at 37°C. Reactions were then separated into supernatant and pellet, and the pellet was retrieved by resuspension in the reaction buffer using a volume equal to that of the reaction. The supernatant and pellet of each sample were separately analyzed by 10% SDS-PAGE gel and visualized by Coomassie blue. After digital scanning, gels were analyzed by FIJI (47) to measure the intensity of binding protein (BP) in the supernatant (S) and pellet (P) fractions. Then, we divided ‘BP in the pellet (P)’ by the ‘total BP (S+P)’ in the reaction to get the ‘fraction of BP in the pellet’.

For all F-actin binding assays, CLIP-170 fragments usually had some minor self-pelleting in PEM50 buffers. In theory, the fraction of self-pelleting might affect the F-actin binding measured, so we always ran a protein-only sample to determine the percentage of self-pelleting. However, we found the self-pelleting in all CLIP-170 proteins and mutants were very similar except for the K70A mutant, which had more severe self-pelleting and so was not used for the follow-up binding assays. To account for self-pelleting behavior, we subtracted the fraction of self-pelleting protein (the ‘fraction of BP in the pellet’ in the BP-only sample) from the ‘fraction of BP in the pellet’ to obtain the ‘fraction of BP bound’, which was then used to calculate the F-actin binding affinity (Figure 1B). The fraction of BP bound for all samples in this work was assessed in this way.

Unless otherwise indicated, the standard reaction buffer was PEM50 (50 mM PIPES, 2 mM MgCl_2_, 1 mM EGTA, pH 6.8) for F-actin binding assays, and PEM buffer (100 mM PIPES, 2 mM MgCl, 1 mMEGTA, pH 6.8) for MT binding assays. The pH values of all buffers were adjusted by KOH. To accommodate the lower salt condition for salt sensitivity assays (Figure 1C), we used a different base buffer (20 mM PIPES, 2 mM MgCl_2_, 1 mM EGTA, pH 6.8), and we used KCl to adjust the salt concentration up to the desired levels. The lowest salt concentration that can be reached after adjusting the pH of the 20 mM PIPES base buffer is 44 mM; 150 mM represents the concentration of salt at physiological conditions (27). All ‘salt concentration’ labels in this study refer to the total potassium concentrations from the KOH and KCl added in the buffer. Protein concentrations were adjusted according to the experiment needs and are indicated in the figure legends.

To estimate the apparent K_D_ from the resulting data (Figure 2 B, C), the binding curves of each CLIP-170 construct were fitted to a biomolecular simple binding equation (with the assumption of a 1:1 binding ratio): Y = B_max_ * X/(K_D_+X), where Y is the fraction of CLIP-170 construct in the pellet, X is the concentration of free F-actin, and B_max_ is maximal achievable binding, which was set to 1. Analysis was performed using OriginPro. Free F-actin was calculated by assuming a 1:1 binding interaction and then subtracting the concentration of CLIP-170 bound (i.e. the concentration of CLIP-170 in the pellet) from the concentration of total F-actin.

### F-actin bundling assays and MT/tubulin competition assays

The F-actin bundling ability of CLIP-170 fragments was assessed by using both low-speed cosedimentation assays and microscopy (Figure 5). CLIP-170 fragment (4 μM) was mixed with F-actin (5 μM) and Alexa-488-phalloidin (0.8 μM) in a 60 μL reaction. 10 μL of the reaction was moved to another tube right after mixing and incubated separately from the remaining 50 μL reaction. The 10 μL reaction was used for the subsequent microscopy assay, and the remaining 50 μL was used for the low-speed cosedimentation assay. All samples were incubated for 25 min at room temperature before being used in the low-speed cosedimentation assay or microscopy.

Competition assays (Figure 6) were performed by incubating Taxol-stabilized MTs (with 10 μM Taxol) or tubulin dimer, CLIP-170 fragment (H1 or H2), and 0.8 μM Alexa-488-labeled-phalloidin for 5 min at room temperature in PEM50. F-actin was added last to compete with the CLIP-170:MT interactions for 10 min before low-speed cosedimentation assay or microscopy, depending on the experiment needs. The reaction volume for the competition assay was 100 μL, and the concentration of each protein is indicated in the figure legend (Figure 6).

For the low-speed cosedimentation assay, reactions were centrifuged at 16,000 × g for 4 min at room temperature. Reactions were then separated into supernatant and pellet. Pellets (bundled F-actin) were retrieved by resuspension in PEM50 (volume equal to that of the reaction). The supernatant and pellet of each sample were separated by 10% SDS-PAGE followed by Coomassie blue stain and digital scan. Gels were analyzed by FIJI (47) to determine the fraction of protein in the pellet, which represents the fraction of F-actin or MT bundled by CLIP-170.

For the microscopy assay, 5 μL samples of the bundling assay (described above) were used for imaging. Images were acquired by a TE2000 inverted microscope (Nikon) with a 60x objective (1.4 N.A.) and a 1.5× optivar. The microscope was equipped with a Hamamatsu CMOS camera, and the software was Nikon NIS-Elements BR 413.04 64-bit.

### Bioinformatic studies and tools

To identify CLIP-170 in a range of vertebrate organisms, the full-length human CLIP-170 sequence was used as the query to perform BLASTp searches against the NCBI Reference Protein databases for the following organisms: Human (taxid: 9606), chicken (taxid:9031), cow (taxid:9913), frogs and toads (taxid:8342), mouse (taxid:10088), lizards (taxid:8504), pigs (taxid:9821), bony fish (taxid:7898), and elephants (taxid:9779). The resulting sequence set contains isoforms of CLIP-170 and its close paralog CLIP-115 (48). Sequences were aligned by ClustalX (49). We then used Jalview (50) to remove redundant sequences, which we defined as those with more than 98% percent identity. The conservation of CLIP-170 in this alignment was mapped onto the two CAP-Gly crystal structures (2E3I and 2E3H (30)) from the Protein Data Bank (PDB) using structure analysis tools in Chimera (51). The electrostatic maps were generated by Coulombic Surface Coloring tool in Chimera with default settings.

### Cell culture and immunofluorescence

COS-7 cells (a gift of Dr. Kevin Vaughan) were grown on 10 mm^2^ glass coverslips (Knittel Glaser) in DMEM with 1% glutamine and 10% fetal bovine serum (Sigma). Cells were incubated with 5% CO_2_ at 37°C. To determine the colocalization between CLIP-170 and F-actin, we transfected cells with N-terminal EGFP-conjugated wild-type CLIP-170 (GFP-CLIP-170), which was controlled by a CMV promoter in pCB6 vector (16). After ~24 hr transfection, cells were fixed with 3% paraformaldehyde (PFA) as described previously (52). After PFA fixation, F-actin was labeled with rhodamine-phalloidin (Cytoskeleton, Inc. PHDR1) for 20 min and followed by 3x PBS wash. Mowiol 4-88 mounting medium (Sigma 475904-M) was used to mount cells on slides. Image acquisition was performed with the same microscope and objective described in the competition assays.

## Acknowledgements

This research was funded by an American Heart Association pre-doctoral fellowship (#17PRE33670896) to YOW, and National Science Foundation grants MCB #1817966 and MCB #1244593 to HVG.

